# The Drosophila Amyloid Precursor Protein homologue mediates neuronal survival and neuro-glial interactions

**DOI:** 10.1101/2020.03.09.983353

**Authors:** Irini A. Kessissoglou, Dominique Langui, Amr Hasan, Maral Maral, Suchetana Bias Dutta, P. Robin Hiesinger, Bassem A. Hassan

## Abstract

The amyloid precursor protein (APP) is a structurally and functionally conserved transmembrane protein whose physiological role in adult brain function and health is still unclear. Because mutations in APP cause familial Alzheimer’s disease, most research focuses on this aspect of APP biology. We investigated the physiological function of APP in the adult brain using the fruit fly *Drosophila melanogaster*, which harbors a single APP homologue called APP Like (APPL). Previous studies have provided evidence for the implication of APPL in neuronal wiring and axonal growth through the Wnt signaling pathway. However, like APP, APPL continues to be expressed in all neurons of the adult brain where its functions and their molecular and cellular underpinnings are unknown. We report that APPL loss of function results in the dysregulation of endolysosomal function, in both neurons and glia, with a notable enlargement of early endosomal compartment in neurons followed by neuronal cell death, the accumulation of dead neurons in the brain during a critical period at a young age and subsequent reduction in lifespan. These defects can be rescued by reduction in the levels of the early endosomal regulator Rab5, indicating a causal role of endosomal function for cell death. Finally, we show that the secreted extracellular domain of APPL is taken up by glia, regulates their endosomal morphology and this is necessary and sufficient for the clearance of neuronal debris in an axotomy model. We propose that the APP proteins represent a novel family of neuro-glial signaling proteins required for adult brain homeostasis.

## Introduction

Early-onset familial Alzheimer’s disease (fAD) is caused by several mutations either in the Amyloid Precursor Protein (APP) or in the Presenilin (PSEN-1 and PSEN-2) genes [1, 2]. APP is a functionally and structurally conserved transmembrane protein, present in both invertebrates like *Caenorhabditis elegans* and *Drosophila melanogaster* [3, 4] and mammals [5, 6, 7]. APP undergoes two competing proteolytic processes; the amyloidogenic processing where it is internalized into endosomes and cleaved by β-secretase and subsequently γ-secretase releasing sAPPβ, the amyloid-β (Aβ) oligomers and APP intracellular domain (AICD), and the non-amyloidogenic processing where APP is cleaved on the cellular membrane by α-secretase and subsequently γ-secretase releasing sAPPα, the P3 domain and AICD [8].

FAD mutations result in the enhancement of the amyloidogenic processing of APP and hence in an increased release of Aβ oligomers, but also, a reduced production of sAPPα [9] and potentially other unknown effects on APP’s physiological function, such as the balance between its intracellular and extracellular activities. The accumulation of Aβ oligomer aggregates is also present in the brain of patients with sporadic Alzheimer’s Disease (AD), forming the Aβ plaques and leading to the hypothesis that Aβ plaques are the main cause of the disease [10]. However, thus far all anti-amyloid treatment, although often successful in reducing the amyloid load, have failed to improve AD symptoms [11]. This raises the need for a better understanding of the physiological function of APP in order to design better future treatment.

*In vitro* loss of function (LOF) studies on human or mouse APP revealed its involvement in a variety of functions related to neuron biology, such as neural stem cell proliferation, differentiation and neurite outgrowth of hippocampal neurons [12]. Moreover, it seems to have a role in synapse formation, as a synaptic adhesion molecule [13]. APP’s conserved intracellular domain interacts with many protein-signaling pathways such as, the JNK to induce cell death [14], X11/JIP to activate cell differentiation [15] and with Fe65 to modulate gene transcription [16].

In *Drosophila melanogaster*, APP-Like (APPL) is the single homologue of the human neuronal APP_695_ sharing 30% homology at the amino acid level [17]. *In vivo* LOF studies have demonstrated that APPL is involved in axonal outgrowth during development [18], axonal transport of vesicles or mitochondria [21, 20], synapse formation at the neuromuscular junction [19] and long-term and working memory formation [22, 23]. Moreover, it has been shown that the sAPPL has a neuroprotective function by rescuing vacuole formation in the brain of neurodegenerative mutant flies through its binding to the full length APPL [24]. Finally, like in mammals, APPL acts as a receptor and interacts with G_0_ proteins, cell adhesion molecules, and intracellular modulators like the dX11/Mint protein, Tip60 and Fe65 [25, 26, 27].

An interesting observation is that most APP LOF studies in a plethora of neuronal processes and molecular mechanisms reveal relatively mild phenotypes with relatively low penetrance. Combined with the fact that neuronal forms of APP are expressed throughout the brain, this suggests that APP is a homeostasis factor required for the brain to develop correctly, remain stable and counteract internal and external perturbations. The nervous system encounters several types of genetic mutations and environmental perturbations that can cause organelle stress, cell death and finally can lead to developmental, age or stress associated disorders. To counteract this, animals have evolved a defense homeostatic signaling system, composed of protein chaperones and transcriptional mechanisms [28] involving both neurons and glial cells such as Astrocytes, Schwann cells and oligodendrocytes [29]. However, the molecules that neurons use to communicate homeostatic signals to glia remain largely unknown.

A major homeostatic cellular mechanism is the endolysosomal recycling and degradation pathway [30]. This pathway ensures that cellular cargo is properly recycled between the membrane and various organelles or degraded to maintain protein homeostasis and cellular health. A study on primary neurons revealed that an APP intracellular binding protein, PAT-1, regulates the number of early endosomes and endocytosis [31]. Recently, two studies revealed that iPSC-derived human neurons with either APP or PSEN1 fAD knock-in mutations show enlarged and defective early endosomes and lysosomes [32, 33]. Therefore, this might suggest a role for APP in the neuronal endolysosomal pathway.

To investigate the *in vivo* role of APP in neuronal homeostasis we used *Drosophila* as a model organism and investigated the consequences of the deletion of its homologue, the *Appl* gene. We report that loss of APPL results in the increased accumulation of apoptotic cells in the brain at a critical young age. We link this accumulation to defects in the endolysosomal pathway in both neurons and glia and show that APPL is required for neuro-glial communication.

## Results

### APPL is required for neuronal survival in young adult flies

To investigate the implication of APPL in brain health of adult flies, we started with quantifying the survival of APPL null flies (*appl*^*d*^) [34] compared to genetic background controls (Canton S) at different stages of their lifespan. As previously reported [24], *appl*^*d*^ flies die significantly earlier than their control counterparts in a sex-independent manner starting at 2-3 weeks of age (Figure 1a). This suggests that loss of APPL compromises survival at an early age. Because *Drosophila* APPL is an exclusively neuronal protein [17], we asked whether neuronal health is compromised in APPL mutants during the first 3 weeks of life. We measured the cell death load in the brain of *appl*^*d*^ and controls at 2, 7, 21 and 45 days of age. To quantify the number of dying cells at any given moment, we stained whole mount brains with Cleaved Drosophila Death caspase protein-1 (Dcp-1), the homologue of human Caspase 3, and manually quantified the Dcp-1 positive cells across the entire brain (Figure1b, b’). In both genotypes, 2 day old flies show significant cell death in their brain due to ongoing brain remodeling [35]. By 7 days of age however, there is a sharp drop in the number of apoptotic cells in controls. In contrast, the drop in apoptosis is significantly reduced in *appl*^*d*^ flies, with an average of 7-8 apoptotic cells per brain at any given time point. Counter staining with the neuronal marker Elav and the glial marker Repo showed that all dying cells detected were neurons (Figure 1b’’, b”‘, b’’’’, b’’’’’). With age, at 21 and 45 days old, both control and *appl*^*d*^ flies show a similar increase in apoptotic cells (Figure 1c). These data suggest that loss of APPL renders neurons particularly sensitive during the first week of life. APPL is only detectable in neurons although some reports have claimed it may be expressed in glia [36]. To test whether APPL expression in neurons is required for their survival at 7 days old, we knocked-down the expression of APPL using the UAS-Gal4 system expressing APPL RNAi only in neurons using the pan-neuronal *nSyb-Gal4* driver. This resulted in significantly more apoptotic neurons in APPL knock-down flies compared to controls, similar to *appl*^*d*^ flies (Figure 1d).

**Figure 1.**
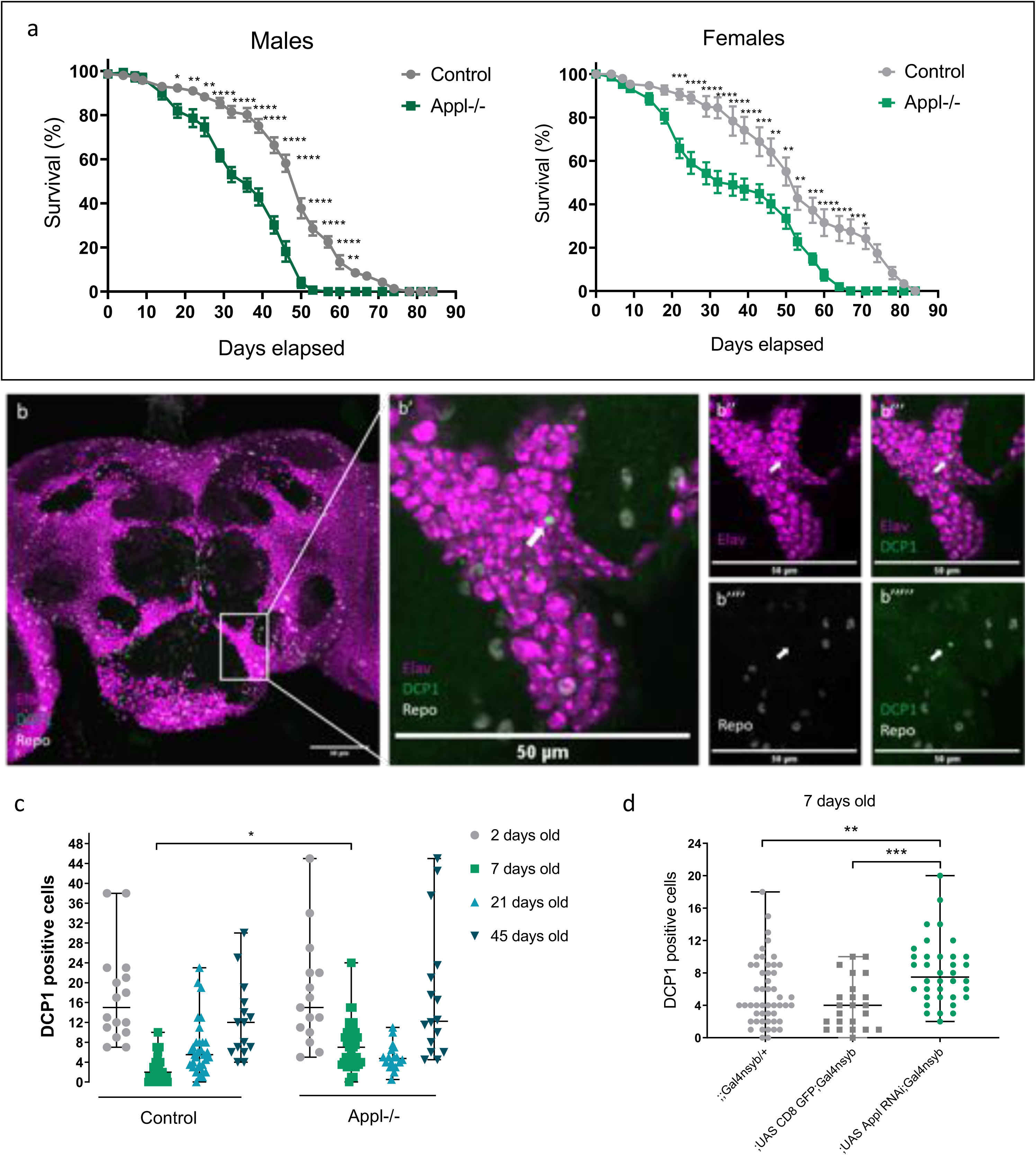
Loss of APPL increases early age mortality rate and apoptotic neuronal death. a) These survival curves represent the lifespan of *appl*^*d*^ (APPL-/-) flies compared to control, Canton S flies. n=10 groups of 15 flies each for every condition (CS males, CS females, APPL-/- males and APPL-/- females). Two-way ANOVA with Tukey’s multiple comparison test df=3, *p<0.05, **p<0.01, ***p<0.0001. b) Confocal sections of an *appl*^*d*^*/Y;;* brain stained with anti-Elav (magenta) to mark the neurons, anti-Cleaved Drosophila Dcp-1 (green) to mark the apoptotic cells and anti-Repo (white) to mark the glial cells. In the higher magnification pictures on the right panels we can notice that the Dcp-1 marked cell (white arrow) colocalises with Elav (b’’-b’’’) but not with Repo (b’’’’-b’’’’’). c) This graph shows the number of apoptotic cells in the central brain of Control and APPL-/- flies at different ages; 2, 7, 21 and 45 days old. Each data point represents the number of apoptotic cells, Dcp-1 positive cells, in a single brain. For the analysis of this data we used Two-way ANOVA with Bonferroni’s multiple comparison test df=3, \**p*=0.027. d) Focusing on the 7 days old time point, which showed significant difference in the previous graph (c), we now knock down the expression of APPL only in neurons using the *yw;UAS APPL RNAi (y+); Gal4nsyb* and find a similar increase in the number of apoptotic cells comparing to the controls: *yw;;Gal4nsyb/+* and *w-;UAS CD8 GFP;Gal4nsyb*. One-way ANOVA: df=2, \*\*\**p=*0.0078, \*\*\**p*=0.0005.

In summary, our data show that loss of APPL in neurons results in excessive neuronal death during the first week of life and a corresponding reduction in life starting 1-2 weeks later. We next asked by what mechanism APPL acts to protect neurons and flies from premature death.

### APPL regulates the size and number of neuronal early endosomes

We have previously shown that APPL is a neuronal modulator of the Wnt PCP pathway for the axonal outgrowth during development. Specifically, loss of APPL sensitizes growing axons to reduction in Wnt-PCP signaling and renders the PCP core protein VanGogh (Vang) haploinsufficient [18]. We started by examining the genetic interaction between *appl* and *vang* by removing one copy of *vang* in the *appl*^*d*^ background and measuring neuronal death at 7 days of age. In contrast to the developmental effect on axon growth, we found no effect on the number of apoptotic neurons (Figure S1) suggesting a different mechanism.

A number of observations suggest a tight link between the APP family of proteins and endolysosomal trafficking. First, both human APP and fly APPL carry highly conserved endocytic motifs in their intracellular domains [37], which interact with proteins involved in endocytosis [31]. Second, APP has been implicated in the regulation of the endocytosis of cell surface receptors [38]. Third, as the endolysosomal pathway is involved in APP’s and APPL’s cleavage by β- and γ-secretases, perturbations of the endolysosomal pathway can have negative repercussions in the proteolytic processing of APP and hence the amount of Aβ produced [39]. Fourth, two recent *in vitro* studies showed evidence for the development of enlarged early endosomes and lysosomes in human iPSCs with various fAD mutations in the *APP* and *PSEN1* genes [32, 33]. We therefore investigated whether loss of APPL causes defects in endolysosomal function in the brain.

We used an acidification-sensing double fluorescent (DF) tag, composed of a pH sensitive GFP (pHluorin) and pH-resistant mCherry fused to a myristoylated residue to track all plasma membrane trafficking (myr-DF) [40]. This probe allows the tracking of the trafficking of membrane cargo through the endolysosomal pathway. In neutral pH vesicles, like early endosomes, the probe will fluoresce in both green and red channels, while in acidic vesicles, such as late endosomes and lysosomes, the GFP signal will be quenched and the probe will fluoresce only in red (Figure 2a). Differences in fluorescence values between controls and mutant would indicate potential defects in endolysosomal trafficking. The myr-DF probe was expressed in all neurons, using the *nSyb-Gal4* driver, in control and *appl*^*d*^ flies (Figure 2c, c’). We focused our imaging and quantifications on two easy to identify neuronal populations; the Kenyon cells of the mushroom body and the Projection Neurons of the antennal lobes (Figure 2b). In live confocal imaging data of 7-day-old flies, due to the diffused green signal of pHLuorin, the probe does not distinguish between early and late endosomes, but permits to measure the quantity and volume of the endolysosomal compartments. This analysis revealed the presence of significantly enlarged endolysosomal compartments in the neurons of *appl*^*d*^ flies compared to those of controls (Figure 2c-d).

**Figure 2.**
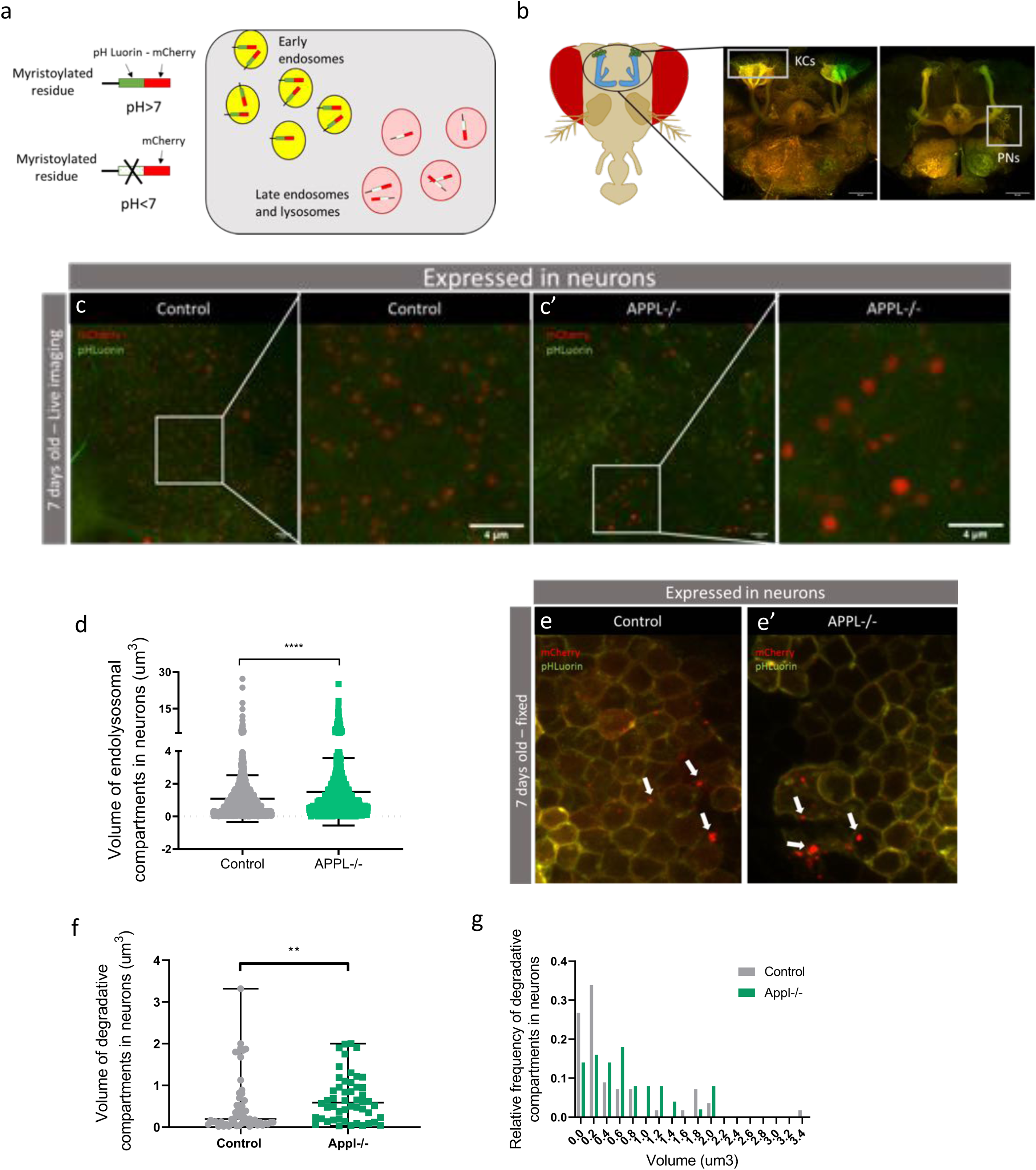
Loss of APPL causes enlarged endolysosomal compartments in neurons. a) Schematic showing the double fluorescent probe composed of a pH-sensitive pHLuorin and pH-resistant mCherry. This probe gets tagged on Myristoylated general plasma membrane proteins. When inside an early endosome it emits a yellow signal and as soon as the protein cargo is going inside an acidic vesicle then it produces a red signal. b) This cartoon represents the head of a fly. The fluorescent images on the right highlight the areas that were imaged: the Kenyon cells (KCs) and Projection neurons (PNs). c-c’)These pictures are from a live snapshot of the KCs of adult, 7 days old, fly brains expressing the double fluorescent probe only in neurons. The left panel and its close-up, show a control: *w*;UAS myr mCherry pH Luorin; nsyb Gal4* fly and the right panel and its close-up, an *appl*^*d*^ mutant: *APPLd; UAS myr mCherry pH Luorin; nsybGal4*. d) This graph shows the quantification of the volume of endolysosomal compartments (um3). n=2 brains per genotype,\*\*\*\**p*<0.0001, Mann-Whitney *post-hoc* test. e-e’) These confocal slices represent the same area of Kenyon cells but this time from a fixed tissue of control: *w*;UAS myr mCherry pH Luorin; nsyb Gal4* and mutant: *APPLd; UAS myr mCherry pH Luorin; nsybGal4* 7 days old fly brains. The white arrows show the acidic degradative compartments. f) The volume of these degradative compartments is significantly higher in *appl* ^*d*^ flies, \*\**p*=0.006, Mann-Whitney *post-hoc* test. g) A histogram of the relative frequency of degradative/acidic compartments in neurons.

Live imaging data could only inform us about the trafficking of the protein cargo to an acidic vesicle, with a pH below 6, but not its degradation inside this vesicle [40]. To quantify the effect of APPL LOF on the degradation of plasma membrane protein cargo we quantified red-only compartments in fixed tissue, where irreversibly damaged pHLuorin leads to the selective loss of green fluorescence [40]. Results from 7 days old fixed brains showed a marginal but not significant increase in the number of degradative compartments between control and *appl*^*d*^ brains (Figure 2e, e’, Figure S2c). However, the volume of degradative compartments in *appl*^*d*^ flies was significantly larger (Figure 2f, g). Together, these analyses evoke the possible enlargement of late-endosome-like vesicles, suggesting a deficit in the regulation of the volume of endolysosomal compartments in *appl*^*d*^ flies.

To further investigate these potential defects at higher resolution, we used Transmission Electron Microscopy (TEM) (Figure 3a, b). Whereas the overall size of the neuronal cell body did not differ between mutants and controls (Figure 3c), we noted the presence of enlarged clear-single membraned endosome-like vesicles with darker content in *appl*^*d*^ neurons (Figure 3a, d). In addition, there were more of them per section than in controls (Figure 3e, f). On the other hand, lysosomal size did not seem to be affected in *appl*^*d*^ flies (Figure 3b, g) and there was a marginal but significant increase in lysosomal number per section (Figure 3h). These data confirm the presence of defects in the neuronal endolysosomal pathway in the absence of APPL, and suggest that these defects arise mostly in endosomes.

**Figure 3.**
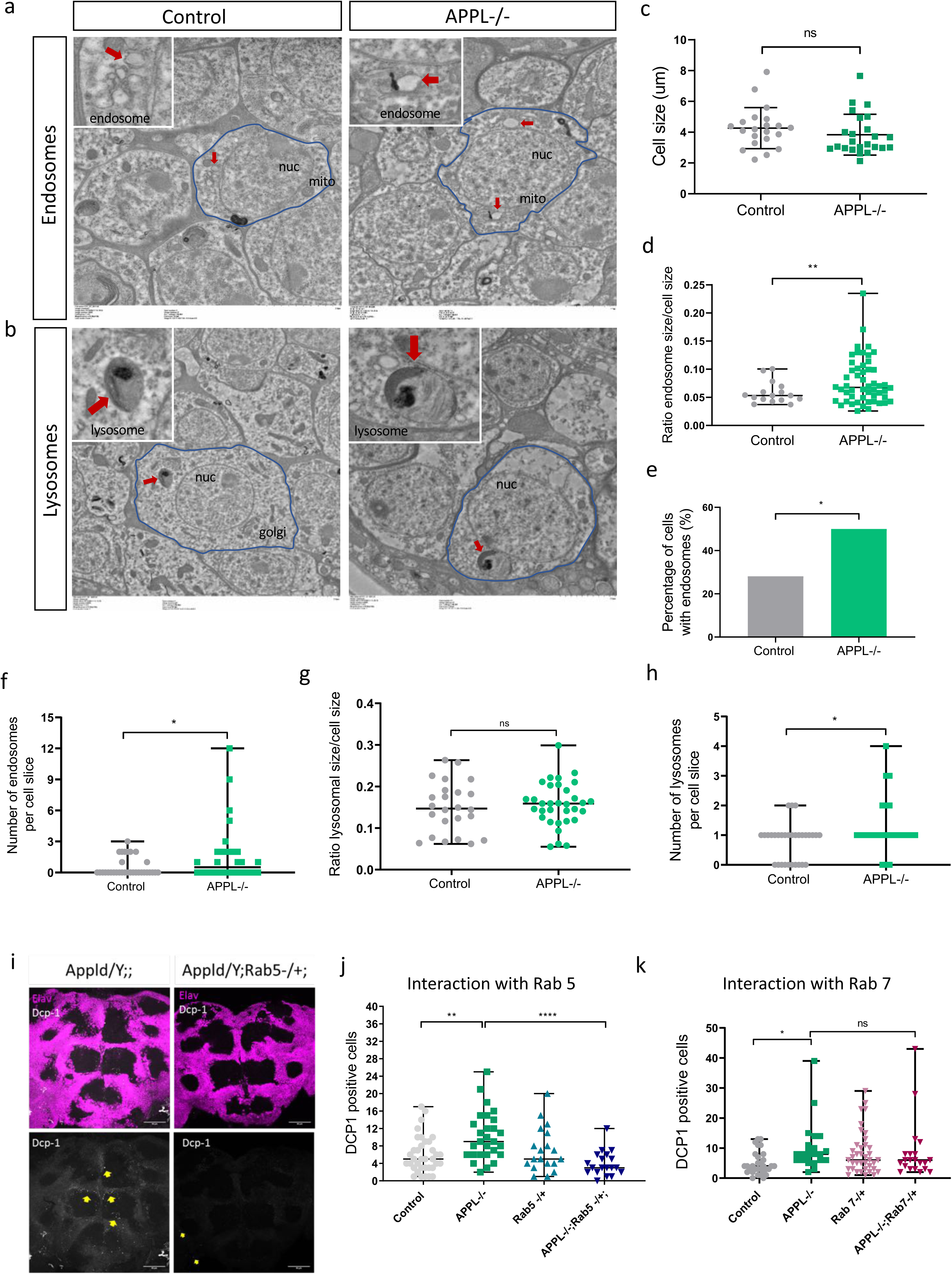
APPL regulates the size of early-endosomes in neurons. a) TEM horizontal sections of the cortical region of a 7 days old fly brain showing neuronal cell bodies (circled in blue) and their organelles. We can observe that there are more and enlarged early-endosome like vacuoles (red arrow) in the brain of APPL-/- flies *w*appl*^*d*^*/Y;;* comparing to control +/+;+/+;+/+. nuc=nucleus, mito=mitochondria b) The size of lysosomes (red arrow) seem to not be affected in APPL-/- flies. c) This graph shows that the cell size is the same between control and APPL-/- flies at 7 days old. d-f)These graphs present the difference in size between the early-endosome like vacuoles seen in APPL-/- and control fly brains and the increased prevalence of endosomes in 7 days old APPL-/- fly brains; n=3 brains per genotype and a total of 32 cells analysed per genotype. Statistical analysis was done using (d) Welch’s t test \*\**p*=0.0080, (e) Binomial test: \**p*=0.0141 and (f) Welch’s t test \**p*=0.0412. g) This graph shows that the size of lysosomes is not affected by the absence of APPL. h) This graph shows that there are significantly more lysosomes per cell slice in APPL-/- flies comparing to Canton S. i) Confocal sections of the central brain of 7 days old APPL-/- flies and APPL-/- flies heterozygous for Rab5, stained with the neuronal marker elav (magenta) and the apoptotic marker dcp-1 (white). Yellow arrows point to the dcp-1 positive cells. j) Quantification of apoptotic cells in the central brain of control *w*/Y;+/+;+/+, w*appl*^*d*^*/Y;;, w*/Y;Rab5 KO/+;* and *w*appl*^*d*^*/Y;Rab5 KO/+;*. Reducing one copy of Rab5 in an APPL-/- background shows a significant reduction in the number of Dcp-1 positive cells observed in APPL-/- flies at 7 days old. One-way ANOVA: df=3, \*\**p*=0.0013 \*\*\*\**p*<0.0001. k) 7 days old APPL-/- flies lacking one copy of Rab7, a late endosome marker, *w*appl*^*d*^*/Y;;Rab7 KO Crispr 3P3RFP/+*, do not show a difference in the number of apoptotic neuronal cell death.

Our data so far suggest a defective accumulation of enlarged endosomes in neurons of *appl*^*d*^ flies. The trafficking of cargo from the membrane to early endosomes is regulated by the Rab5 GTPase. To investigate whether defects in early endosomes cause the increase in the number of dying neurons in *appl* mutant brains, we removed one copy of the *rab5* gene in an *appl* null background. This completely rescued the neuronal cell death phenotype at 7 days of age back to control levels (Figure 3i-j). Moreover, reducing a copy of *rab5* rescued the early death rate and extended the lifespan of the *appl* null flies (Figure S4a). We conclude that reducing the trafficking to early-endosomes in an *appl* null condition re-equilibrates the system and rescues the functioning of the endolysosomal pathway.

To test whether the rescue effect is specific to the early endosomal stage, we removed a copy of the gene encoding the late endosomal marker Rab7 in *appl*^*d*^ mutant background. In contrast to reduction of Rab5 levels, this failed to rescue the number of apoptotic cells found in the brains of 7 days old *appl*^*d*^ flies (Figure 3k), and indeed significantly worsened the lifespan of the flies relative to controls (Figure S4b), consistent with a role for Rab7 itself in neurodegeneration [51].

Our observations suggest that in the absence of APPL, neurons accumulate enlarged vacuole-like endosomal compartments, possibly due to the dysregulation of early endosomes, resulting in neuronal death in the young adult brain and an eventual shortening of lifespan. What is intriguing, however, is why these dying neurons accumulate to a sufficient level as to be detectable instead of being cleared by glial cells. We therefore wondered whether the absence of APPL may be causing a problem with glial clearance.

### The extracellular domain of APPL is secreted by neurons and taken up by glia

APPL is a transmembrane protein that is cleaved resulting in a secreted form, APPLS. To explore the expression and secretion pattern of APPL, we generated a double-tagged form of APPL (dT-APPL) with GFP intracellularly (C-terminally) and mCherry extracellularly (N-terminally) (Figure 4a). To study the distribution and spread of APPLS, we expressed dT-APPL strictly in the retina and imaged the entire brain at different stages of pupal development and in the adult. Whereas the intracellular part of the *appl* protein (GFP), remained inside photoreceptors, APPLS (mCherry), gradually spread throughout the whole brain starting from 80H after puparium formation and remained so in adults (Figure S6a-c). Moreover, APPLS was taken up by glia (Figure S6c). To ascertain that glial uptake of APPLS was not a consequence of APPL overexpression in the presence of the endogenous protein, we repeated this experiment by expressing the dT-APPL in neurons of *appl* null flies. Again, while the intracellular part of APPL remained in neurons, APPLS was localized both in neurons and in glia (Figure 4b-b’’’). Therefore, these data suggest a non-cell autonomous function for APPLS, in glia.

**Figure 4.**
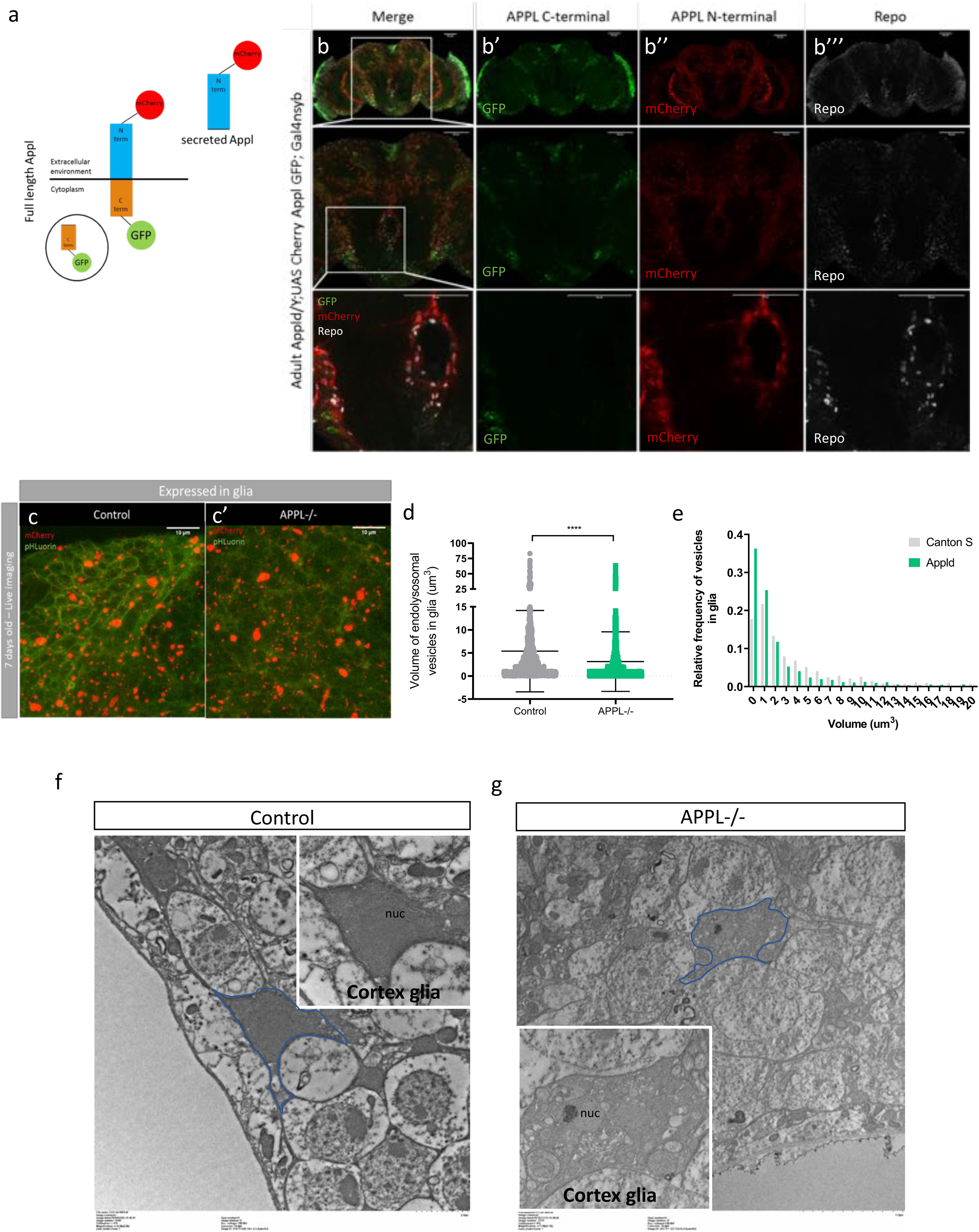
SAPPL interacts with glia and affects their endolysosomal network. a) Schematic representation of the double-tagged APPL construct with GFP on the intracellular part and mCherry on the extracellular part. b-b’’’) Confocal sections of an adult APPL null fly brain expressing the double-tagged construct *Appldw*/Y; UAS CherryApplGFP / +; nsyb Gal4* in all neurons. In green we see the intracellular domain C-terminus of APPL and in red the secreted N-terminus of APPL. This brain is also stained with anti-Repo to mark glia (white). On the close-up panels (last row) we can clearly observe the colocalisation between SAPPL and the glial marker, Repo. c-c’) These pictures are from a live snapshot of glial cells, around Kenyon cell bodies, of 7 days old fly brains expressing the double fluorescent probe only in glia. The left panel shows a control: *w*; UAS myr mCherry pH Luorin; repoGal4* fly and the right panel the mutant: *Appldw*; UAS myr mCherry pH Luorin; repoGal4*. d, e) This graph and histogram represent the quantification of the volume of endolysosomal compartments in glia. They show that the endolysosomal compartments are smaller in glia of APPL-/- flies, \*\*\**p*=0.0009, Mann-Whitney *post-hoc* test. f, g) TEM horizontal sections of the cortical region of a 7 days old fly brain showing neuronal cell bodies and cortical glia (circled in blue) between them. We can observe that the distribution of cortical glia in the brain of APPL-/- flies is abnormal, they have a strange shape and many vesicles, comparing to the control.

### APPL regulates glial endolysosomal volume and debris degradation function

Considering the involvement of APPL in the regulation of the size of endosomes in neurons, we asked whether APPLS may play a similar role in glia. We expressed the myr-DF probe specifically in glia and performed live imaging of 7-day-old control and *appl*^*d*^ brains. In contrast to neurons, the endolysosomal compartments of glia had a reduced volume compared to the controls (Figure 4c, c’, d), with no significant effects on their numbers (Figure S5a). The volume and number of degradative compartments analysed from fixed data was not affected by the absence of APPL (Figure S5b-e). TEM analysis however revealed strong glial disruptions. In control brains, cortex glia were intact and their extensions occupied the spaces between neuronal cell bodies (Figure 4f). In contrast, the distribution of cortex glia between neuronal cell bodies in APPL null brains was irregular, and they showed cytoplasmic blebbing, suggesting these glia were either unhealthy or dysfunctional (Figure 4g). These data suggest the exciting possibility that APPLS may act as a neuronal signal to regulate endolysosomal trafficking in glia. Studies on mouse brain lesion models showed increased levels of alpha-secretase (ADAM-17 and ADAM-10) in reactive astrocytes 7 days post-lesion [41]. In *Drosophila*, using a model of axonal ablation of olfactory receptor neurons (ORNs) Kato and colleagues showed that glia lose their ability to react to axonal lesions within 10 days after injury [42]. Therefore, taking into consideration these findings and our data showing a role of APPL during the first week of adulthood in the fly brain and its transfer from neurons to glia, we asked if APPL is required for glia to clear neuronal debris.

To investigate this, we labelled ORNs with GFP in control and *appl*^*d*^ flies and used the model of antennal ablation [43] (Figure 5a). After ablating both antennae of 5 days old flies we dissected their brain and imaged ORN axonal debris (GFP, green) in the antennal lobes of the adult fly brain. In control brains, axonal debris were almost completely cleared by 5 days after ablation. In contrast, loss of APPL caused a significant reduction in the clearance of the degenerative axons by glia in 5 days post-ablation (Figure 5b-f). This defect was rescued by re-expressing, in an *appl* null background, either full length APPL or only APPLS specifically in ORNs (Figure 6a-c). To test the extent of the delay in clearance, we examined control and *appl* null brains at 8 days post ablation, and found that axonal debris still persist in *appl* mutants at this late stage (Figure 6d). Therefore, APPL is a neuronal signal required in glia to regulate their ability to clear neuronal debris.

**Figure 5.**
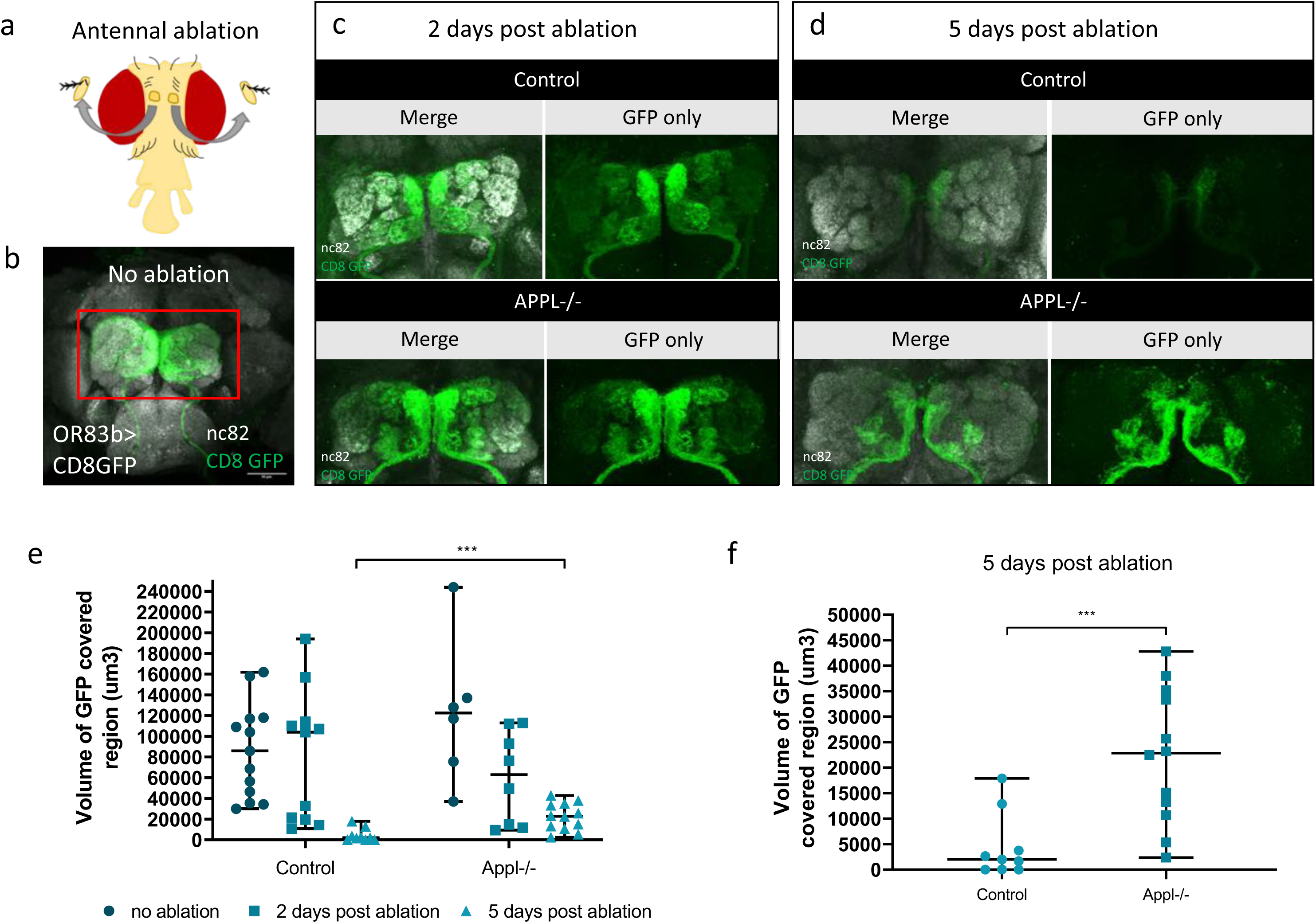
APPL null flies show defective clearance of degenerating axons. a) Schematic presenting the head of a fly after antennal ablation. b-d) Confocal images of GFP-labelled olfactory receptor neuronal axons at the antennal lobes of control: *w*/Y;OR83bGal4 UAS CD8 GFP/+;* and APPL-/-: *Appldw*/Y;OR83bGal4 UAS CD8 GFP/+;* flies before antennal ablation, 2 (c) and 5 (d) days after antennal ablation. The left panels of every section are also stained with nc82 to mark the neuropil. e) Quantification of volume of GFP covered region in the OR83b innervating glomeruli before, 2 and 5 days post ablation, in control and APPL-/- flies. f) We can observe that at 5 days post ablation the volume of GFP covered region of axonal debris remaining in the APPL-/- brains is significantly higher comparing to the control, \*\*\**p*=0.0009, Mann-Whitney *post-hoc* test.

**Figure 6.**
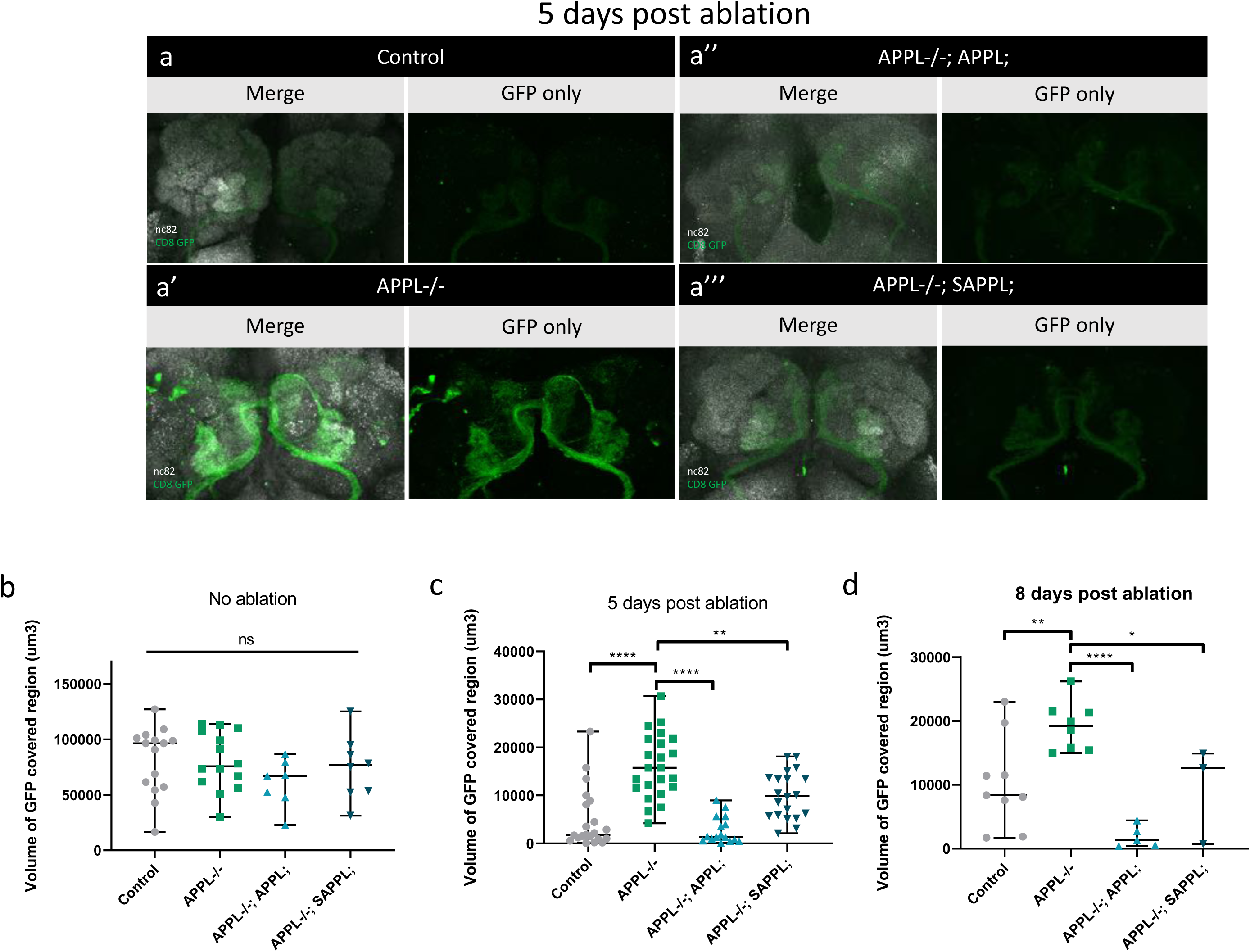
Expression of flAPPL and SAPPL rescues glial clearance of axonal debris. a-a’’’) Confocal images of GFP-labelled olfactory receptor neuronal axons at the antennal lobes at 5 days post antennal ablation. This data show the rescue of a defective glial clearance of axonal debris, seen in APPL-/- flies, when we express the UAS APPL (a’’) and UAS SAPPL (a’’’). The left panels of every section are also stained with nc82 to mark the neuropil. b-d) Quantification of volume of GFP covered region (um3) in the OR83b innervating glomeruli before and at 5 (c) and 8 (d) days post ablation, in control, APPL-/- and the rescue flies: *Appldw*; UAS APPL/OR83bGal4GFP;* and *Appldw*; UAS APPLS/OR83bGal4GFP;.* c) We can observe that at 5 days post ablation, expressing APPL and APPLS in an APPL-/- background are able to significantly rescue the defective glial clearance. One-way ANOVA df=5, \*\**p*=0.0019 \*\*\*\**p*<0.0001. d) This phenotype seems to have a similar pattern at 8 days post ablation. One-way ANOVA df=3, \**p*=0,04 \*\**p*=0.0099 \*\*\*\**p*<0.0001

## Discussion

In this study, we took advantage of *Drosophila melanogaster* to investigate and unravel the physiological function of APPL, the single fly homologue of the human APP, in the adult brain. Our key findings are (1) that APPL is required for neuronal survival during a critical period of early life, (2) regulates the size of endolysosomal vesicles in neurons and glia, and (3) that secreted APPL is taken up by glial cells to enable the clearance of neuronal debris.

### APPL is required for adult brain homeostasis through the endolysosomal pathway

A homeostatic signaling system is composed of a set point, a feedback control, sensors and an error signal. The error signal activates homeostatic effectors to drive compensatory alterations in the process being studied [44]. We propose a model (Figure 7) whereby the presence of APPL and its cleaved forms maintain the physiological flow of vesicular trafficking, either for degradation or for recycling, through the endolysosomal network in neurons. Simultaneously, in case of a system failure, a particular stress or an acute injury, there is increased release of APPLS, the error signal, activating degradation in glial cells, the homeostatic effector, to reset the system to its baseline.

**Figure 7.**
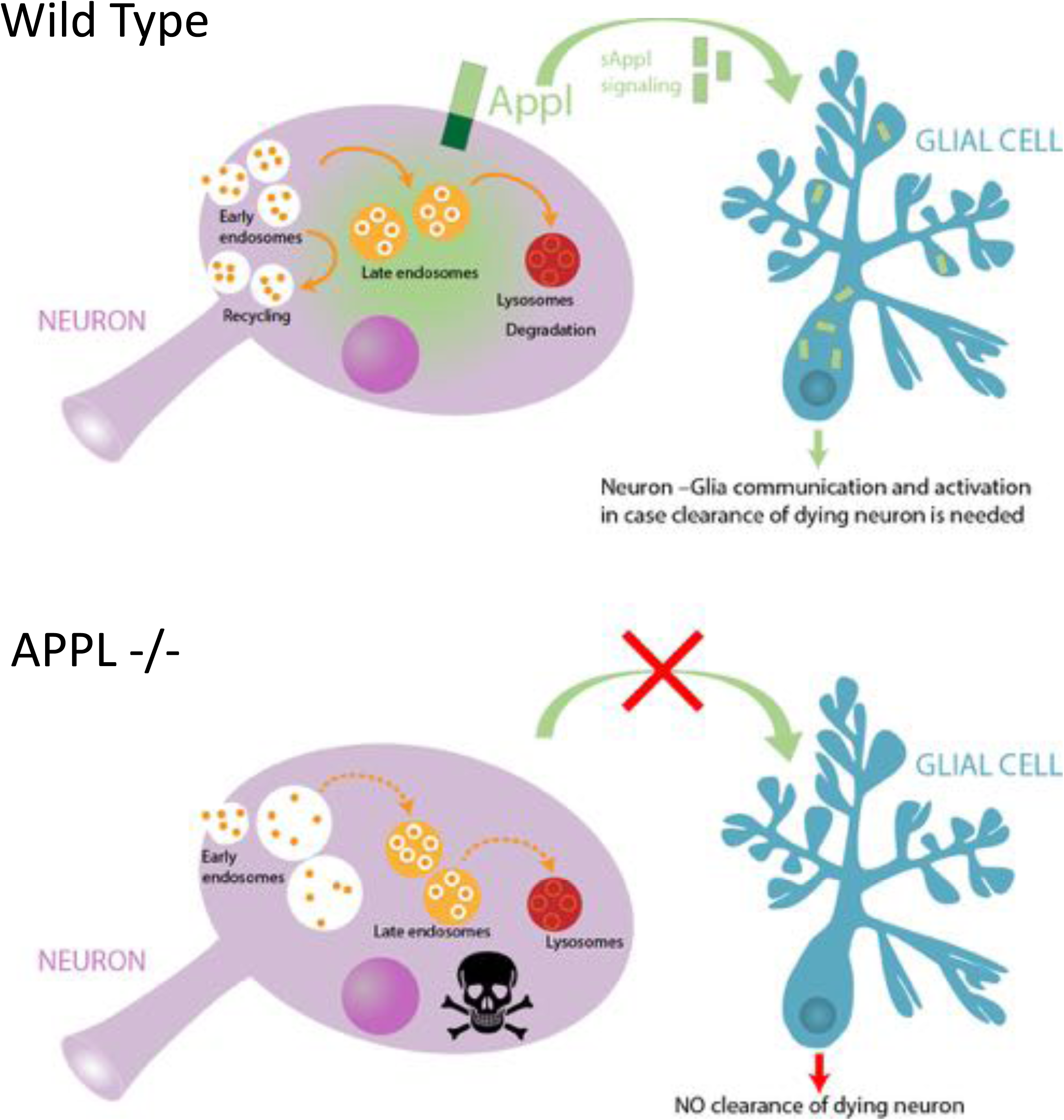
Working model.

It has been observed that *appl* null flies have a shorter lifespan and develop large neurodegenerative vacuoles in their brain by 30 days old [24]. In the present study, we demonstrate that the brain of *appl* null flies shows signs of dysfunctional homeostasis from a much younger age of 7 days old, resulting in a significantly increased number of apoptotic neurons and a significantly increased death rate from 20 days old.

Studies on Down syndrome (Trisomy 21), representing cases of elevated expression of APP, AD patient fibroblasts, AD mouse models and recent studies using patients iPSCs have all shown evidence of a defective endolysosomal network [45, 32, 33]. In particular, neurons derived from AD patient iPSCs show that fAD mutations in APP or PSEN1 as well as knockout of APP all cause alterations in the endo-lysosomal vesicle size and functionality. Some of the toxic effects on endolysosomal trafficking have been attributed not to amyloid accumulation but rather to the potential toxicity of the sAPPβ and/or APP β C-terminal fragment (APPβCTF), while a wealth of literature suggests that full length APP and sAPPα are neuroprotective [46, 24].

### APPL as a neuronal inducer of glial activity

Glial cells are the key immune responders of the brain that maintain neuronal homeostasis through neurotrophic mechanisms and by clearing degenerating neurons. Our data show that neuronal expression of APPL is necessary and sufficient to activate glial clearance of neuronal debris, and that glia take up neuronally released SAPPL. It has also previously been shown that acute injury of the adult brain elicited an increased expression of APPL at and near the site of injury [47]. Interestingly, a recent study using iPSCs derived astrocytes with APP KO and fAD mutations revealed that loss of full-length APP (flAPP) impairs cholesterol metabolism and the ability of astrocytes to clear Aβ protein aggregates [48]. Moreover, upregulation of APP expression in neurons and α-secretase expression in reactive astrocytes was observed after the denervation of the mouse dentate gyrus[41]. Together these observations indicate that the expression and proteolytic processing of APP are part of a neuro-glial signaling system responsible for monitoring brain health and activating glial responses to neuronal injury. Further future work will be needed to describe how exactly secreted APP fragments are taken up by glia and what cellular and molecular components they interact with and modify within glial cells to mediate appropriate levels of glial activation.

### Implications for neurodegeneration

Our findings that the complete loss of the *Drosophila* APP homologue causes deficits in the endolysosomal pathway, in neuron-induced glial clearance of debris and in neuronal death and organismal lifespan strongly suggest that, in the adult brain, the physiological function of full-length APP and the consequences of fAD mutations are mechanistically related to one another. Furthermore, the fact that neuronal death and defective neuronal endosomes are observed very early in life of *appl* mutant flies further supports the notion that significant deficits exist in the AD brain long before any clinical symptoms appear. This may suggest that examining the size and/or function of the early endosome may identify risk for future neurodegeneration and offer future treatment pathways.

## Materials and Methods

### Fly Stocks and Husbandry

**Figure 1** <u>Controls</u>: Canton S (*+/+;+/+;+/+*), *yw*;;nsybGal4, w*;UAS CD8 GFP;, yw; + / +; + / +* kindly given by the lab of T. Preat. <u>Appl-/-:</u> *Appl*^*d*^*w*;+/+;+/+* kindly given by the lab of J-M. Dura and *y1 sc* v1; P{TRiP.HMS01931}attP40; + / +* (UAS Appl RNAi with y+ as a marker) kindly given by the lab of T. Preat.

**Figure 2** <u>Control</u>: *w*;UAS myr mCherry-pHLuorin;, yw*;;nsybGal4.* <u>Appl-/-:</u> *Appl*^*d*^*w*; UAS myr mCherry pH Luorin; nsybGal4* kindly given by the lab of R. Hiesinger.

**Figure 3** <u>Controls</u>: Canton S (*+/+;+/+;+/+*), *w*/Y;+/+;+/+.* <u>Appl-/-</u>: *Appl*^*d*^*w*;+/+;+/+.* <u>Rab5-/+</u>: *w*;Rab5 KO/CyO;* and <u>Rab7-/+</u>: *w*;;Rab7 KO Crispr 3P3RFP/TM6B* kindly given by the lab of R. Hiesinger.

**Figure 4** <u>Control</u>: *w*;UAS myr mCherry-pHLuorin;, w*;;repoGal4* kindly given by V. Auld lab. Canton S (+/+;+/+;+/+). <u>Appl-/-:</u> Appldw*;+/+;+/+. <u>Double fluorescent construct:</u> *Appldw*hsflp / FM7C Df GmR YFP; UAS CherryApplGFP / CyO;* (created in the lab), *yw*;;nsybGal4*. <u>Appl-/-:</u> *Appl*^*d*^*w*; UAS myr mCherry pH Luorin; repoGal4*.

**Figure 5** <u>Control</u>: ;*OR83bGal4 UAS CD8 GFP;* kindly given by the lab of I. Grunwald. <u>Appl-/-</u>: *Appldw*/Y;OR83bGal4 UAS CD8 GFP/+;*.

**Figure 6** <u>Rescue experiment flies</u>: *Appldw*; UAS APPL/OR83bGal4GFP;* and *Appldw*; UAS APPLS/OR83bGal4GFP;.*

**Figure S1** <u>Control</u>: *w*/Y;+/+;+/+* <u>Appl-/-</u>: *w*appl*^*d*^*/Y;;*, and <u>Vang-/+:</u> *appl*^*d*^*w*/Y;Vang-/+;*.

**Figure S6** <u>dT expressed specifically in the retina</u>: *;UAS-mCherry-APPL-GFP/lexAop-CD4tdGFP; GMR-Gal4/Repo-lexA* kindly provided by the lab of R. Hiesinger.

All stocks were maintained using standard rich food at 21°C and all crosses and experiments were conducted at 25°C on a 15hr:9hr light:dark cycle at constant humidity.

### Lifespan experiments

For the lifespan experiment, eclosing adults were collected under CO_2_-induced anaesthesia, over a 12hr period, and were left to mate for 48hrs before sorting them into single sexes. After sorting, they were housed at a density of 15 flies per vial. Throughout the lifespan, flies were kept in a humidified, temperature-controlled, incubator with 15hr:9hr light:dark cycle at 25 °C on a standard, sucrose yeast corn and agar, media. Finally, they were transferred into new food and scored for death every 2-3 days throughout adult life [49].

### Immunochemistry

Adult brains were dissected in phosphate buffered saline (PBS) and fixed in 3.7% formaldehyde in PBT (PBS+Triton 0.3%) for 15min. The samples were subsequently rinsed four times for 0’, 5’, 15’ and 30’ in PBT 0.3% and blocked in 1% BSA for at least 1 hour. Following these steps, the brains were incubated with the primary antibody diluted in 1% BSA overnight at 4°C. Then the samples were rinsed four times for 0’, 5’, 15’ and 30’ in PBT 0.3% and were subsequently incubated with the appropriate fluorescent secondary antibodies in dark for 2 hours at room temperature. Finally, after four rinses with PBT 0.3% the brains were placed in PBS and mounted on a polarised slide using Vectashield (Vector labs) as the mounting medium.

The mounted fixed brains were imaged on an Olympus 1200 confocal microscope equipped with the following emission filters: 490-540 nm, 575-620 nm and 665-755 nm.

The following antibodies were used: rabbit anti-cleaved Drosophila Dcp-1 (Cell Signalling, 1:100), rat anti-elav (Hybridoma bank, 1:100), mouse anti-repo (Hybridoma bank, 1:10) and mouse anti-nc82 (Hybridoma bank, 1:100).

### Transmission Electron Microscopy

First, we cut 7 days old *Drosophila* adult heads and fixed them in 2% glutaraldehyde +2% PFA+ 1mM CaCl2 in 0.1M sodium cacodylate buffer, pH 7.4, for 1hour at room temperature (RT). Following three rinses with Na-cacodylate buffer, we post-fixed samples with 1% osmium tetroxide in the same 0.1M sodium cacodylate buffer for 1h at RT.Then we dehydrated them in a graded series of ethanol solutions (75, 80, 90 and 100%, 10 min each). Final dehydration was performed twice in 100% acetone for 20 min. Subsequently, we infiltrated samples with Epon 812 (epoxy resin) in two steps: 1 night at +4°C in a 1:1 mixture of Epon and acetone in an airtight container and 2h at RT in pure Epon. Finally, we placed samples in molds with fresh resin and cured them in a dry oven at 60°C for 48h.

Blocs were cut in 1 µm semi-thin sections with an ultramicrotome EM UC7 (Leica). Sections were stained with 1% toluidine in borax buffer 0.1M. Then we cut ultra-thin sections (∼ 70 nm thick) and collected them on copper grid (Electron Microscopy Science). They were contrasted with Reynolds lead citrate for 7 min. Observations were made with a Hitachi HT 7700 electron microscope operating at 70 kV. Electron micrographs were taken using an integrated AMT XR41-B camera (2048×2048 pixels).

### Adult brain culture and live imaging

Adult brains were dissected in cold Schneider’s *Drosophila* Medium and mounted, posterior side up, in the culture chambers perfused with culture medium and 0.4% dialyzed low-melting agarose [50]. Live imaging was performed at room temperature using a Leica TCS SP8 X confocal microscope with a resonant scanner, using 63X water objective (+3.3 zoom). White laser excitation was set to 488 nm for pHLuorin and 587 nm for mCherry signal acquisitions [40].

### Quantification and statistical analysis

Imaging data were processed and presented using ImageJ (National Institute of Health). Image J was also used for manual quantification of the apoptotic, dcp-1 positive cells slide by slide throughout the z-stack and for selecting regions of interest using the “ROI Manager” function. For the endolysosomal compartments analysis we used the IMARIS software (Bitplane), for both live and fixed images. To quantify the number and volume of the endolysosomal compartments we used the Surface function, enabling the “Split touching objects” mode and keeping the same intensity threshold across samples and conditions. In the fixed images, to distinguish the red, acidic, compartments from the endosomes and quantify them, we used the “Spot colocalize” function. To measure the volume of the ones non-colocalizing, we used the Surface function enabling the “Split touching objects” mode. Finally, the IMARIS software (Bitplane) and, more specifically, the Surface function was also used to quantify the volume of remaining GFP expressing axonal debris in the antennal ablation experiment, again using the same intensity threshold across samples and conditions. Graphs were generated and statistical analysis was conducted using GraphPad Prism 8.

### Olfactory receptor injury protocol

For the antennal ablation experiment we used *;OR83b Gal4 UAS CD8 GFP;* flies, expressing GFP in most of the olfactory receptor neurons, and crossed them with control and *appl*^*d*^ background flies. The progeny of these crosses was collected daily and, after selecting the right genotype, we ablated both antennae of 5 days old flies using finely sharpened tweezers. Then we dissected the adult brains at 2, 5 and 8 days post ablation and followed the immunostaining procedure, as previously described, in dark. We used anti-nc82 as the neuropil antibody in order to better visualise the antennal lobe glomeruli of the adult brain and focus our quantification of the endogenously expressed GFP covered region accordingly.

## Acknowledgments

We thank the Bloomington stock center (NIH P40OD018537) for providing flies used in this study. We thank all members of the Hassan and Hiesinger labs for support and valuable comments.

## Author contributions

I.A.K. and B.A.H conceived the study, designed the experiments and wrote the manuscript. B.A.H. and P.R.H. supervised the work. I.A.K., D.L., A.H., M.M. and S.B.D. conducted all experiments. I.A.K. performed all data analysis.

## Abbreviations

APP: Amyloid Precursor Protein
APPL: Amyloid Precursor Protein Like
AD: Alzheimer’s disease
fAD: Familial Alzheimer’s disease
PSEN-1/2: Presenilin ½
AICD: APP intracellular domain
Aβ: amyloid β
SAPP: secreted APP
flAPP: full length APP
APPβCTF: APP β C-terminal fragment
JNK: c-Jun N-terminal kinase
PAT-1: Protein interacting with APP tail 1
iPSC: Induced pluripotent stem cell
Dcp-1: Death caspase-1
Vang: VanGogh
DF: double fluorescent
GFP: Green fluorescent protein
Myr-DF: myristoylated double fluorescent
TEM: Transmission Electron Microscopy
ADAM: A Disintegrin and Metalloprotease
ORN: Olfactory Receptor Neuron
PBS: Phosphate buffered saline
RT: room temperature

## Supplementary

**Figure S1.**
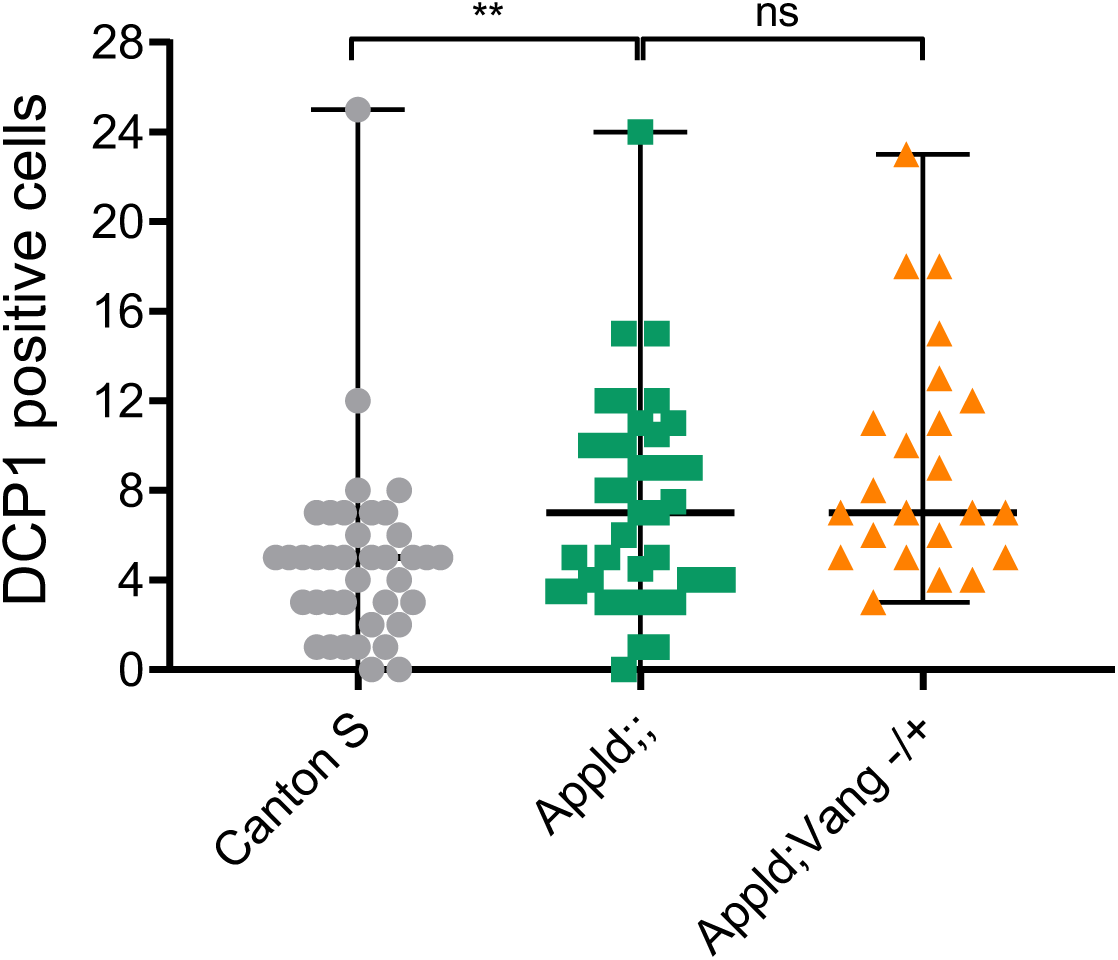
APPL does not seem to interact with the Wnt PCP pathway to maintain neuronal health. Quantification of apoptotic cells in the central brain of control, *w*/Y;+/+;+/+, w*appl*^*d*^*/Y;;*, and *appl*^*d*^*w*/Y;Vang-/+;*. Reducing one copy of Vang, a key member of the Wnt PCP pathway, in an APPL-/- background has no effect on the accumulation of apoptotic cells.

**Figure S2.**
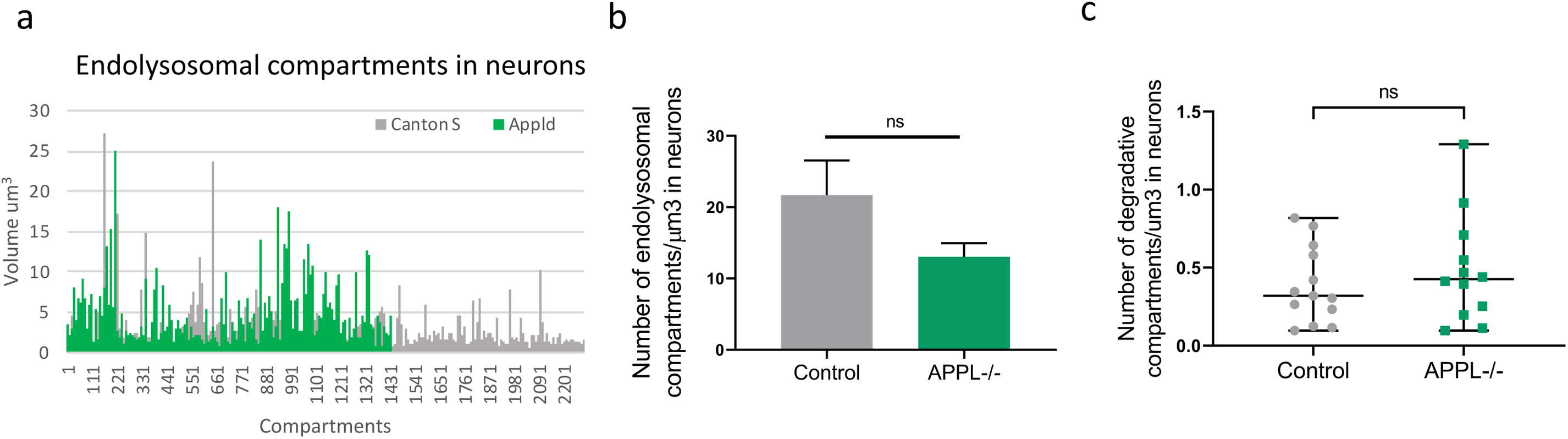
Loss of APPL causes enlarged endolysosomal compartments in neurons. a)Histogram presenting the volume of each endolysosomal compartment in neurons of control, *w*;UAS myr mCherry pH Luorin; nsyb Gal4* fly, and *appl*^*d*^ mutant: *APPLd; UAS myr mCherry pH Luorin; nsybGal4*, flies. b) This graph shows the quantification of the number of endolysosomal compartments. c) This graph shows that the number of degradative compartments/um3 is not significantly affected by the absence of APPL, every dot corresponds to a brain.

**Figure S3.**
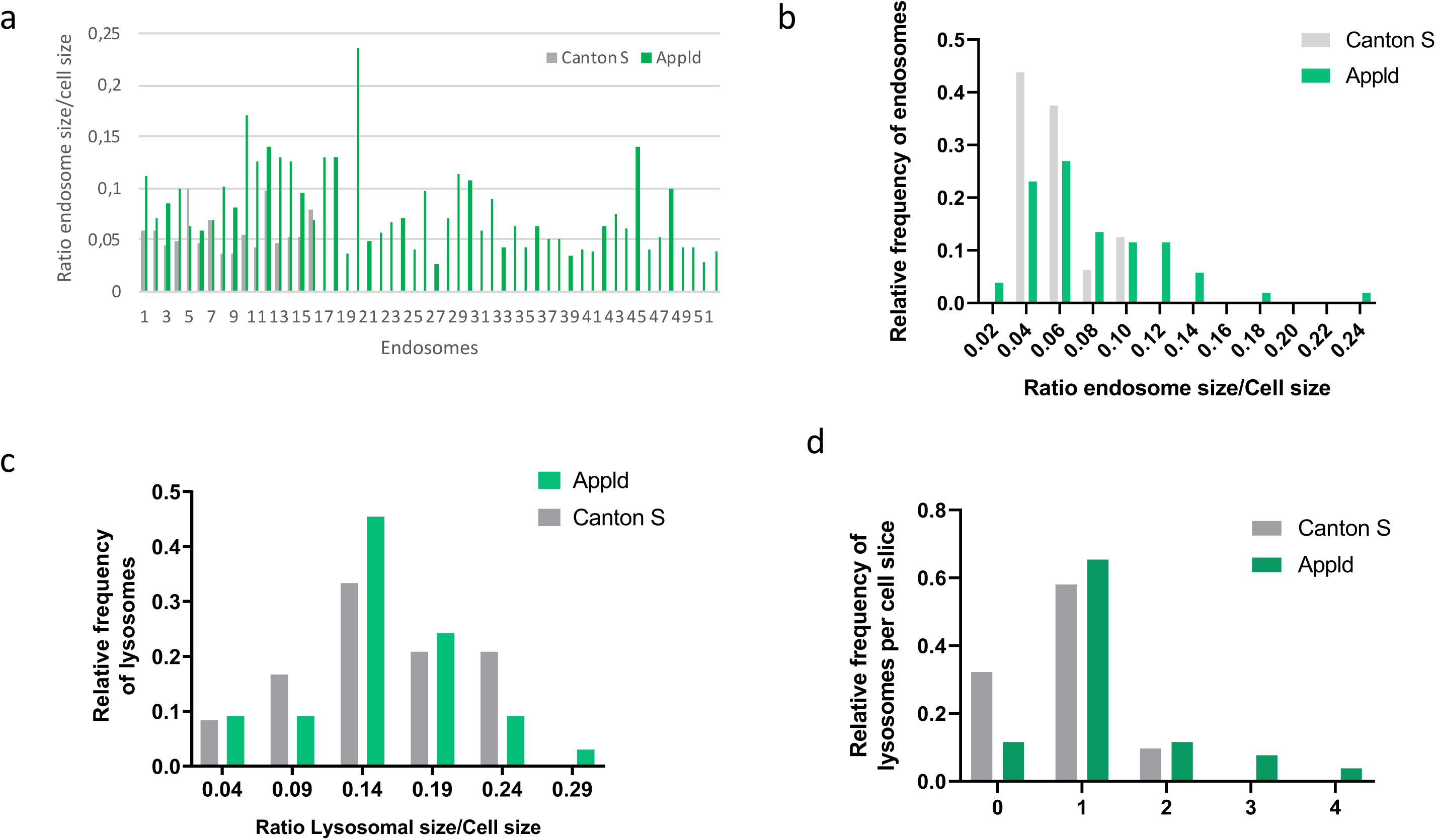
APPL regulates the size of early-endosomes in neurons. a) Histogram presenting the volume of each early-endosome-like vesicle in neurons of control, +/+;+/+;+/+ and APPL-/- flies, *w*appl*^*d*^*/Y;;* flies. b) This histogram represents the relative frequency of early-endosome-like vesicles and their size in APPL-/- flies, *w*appl*^*d*^*/Y;;*, comparing to control, +/+;+/+;+/+. c) This histogram represents the relative frequency of lysosomes and their size in APPL-/- flies, *w*appl*^*d*^*/Y;;*, comparing to control, +/+;+/+;+/+. d) Histogram representing the relative frequency of lysosomes per cell slice in APPL-/- flies, *w*appl*^*d*^*/Y;;*, comparing to control, +/+;+/+;+/+ fly brains.

**Figure S4.**
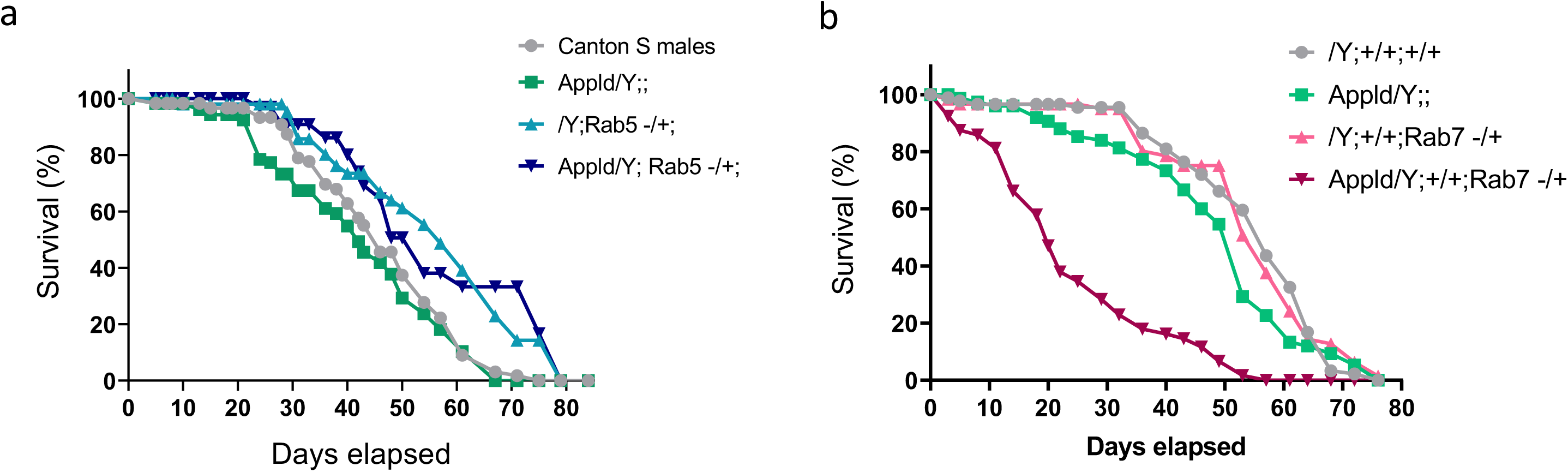
Effects of endolysosomal alterations in lifespan. a-b) Lifespan analysis of control and *appl*^*d*^ flies lacking one copy of Rab5 (a) and *appl*^*d*^ flies lacking one copy of Rab7 (b). These survival curves reveal that reducing one copy of Rab5 in an *appl*^*d*^ background can rescue the early death rate seen in *appl*^*d*^ flies and increase the overall lifespan. Although, reducing one copy of Rab7 in an *appl*^*d*^ background increases significantly the death rate starting from an even earlier age and reducing the overall lifespan of *appl*^*d*^ flies.

**Figure S5.**
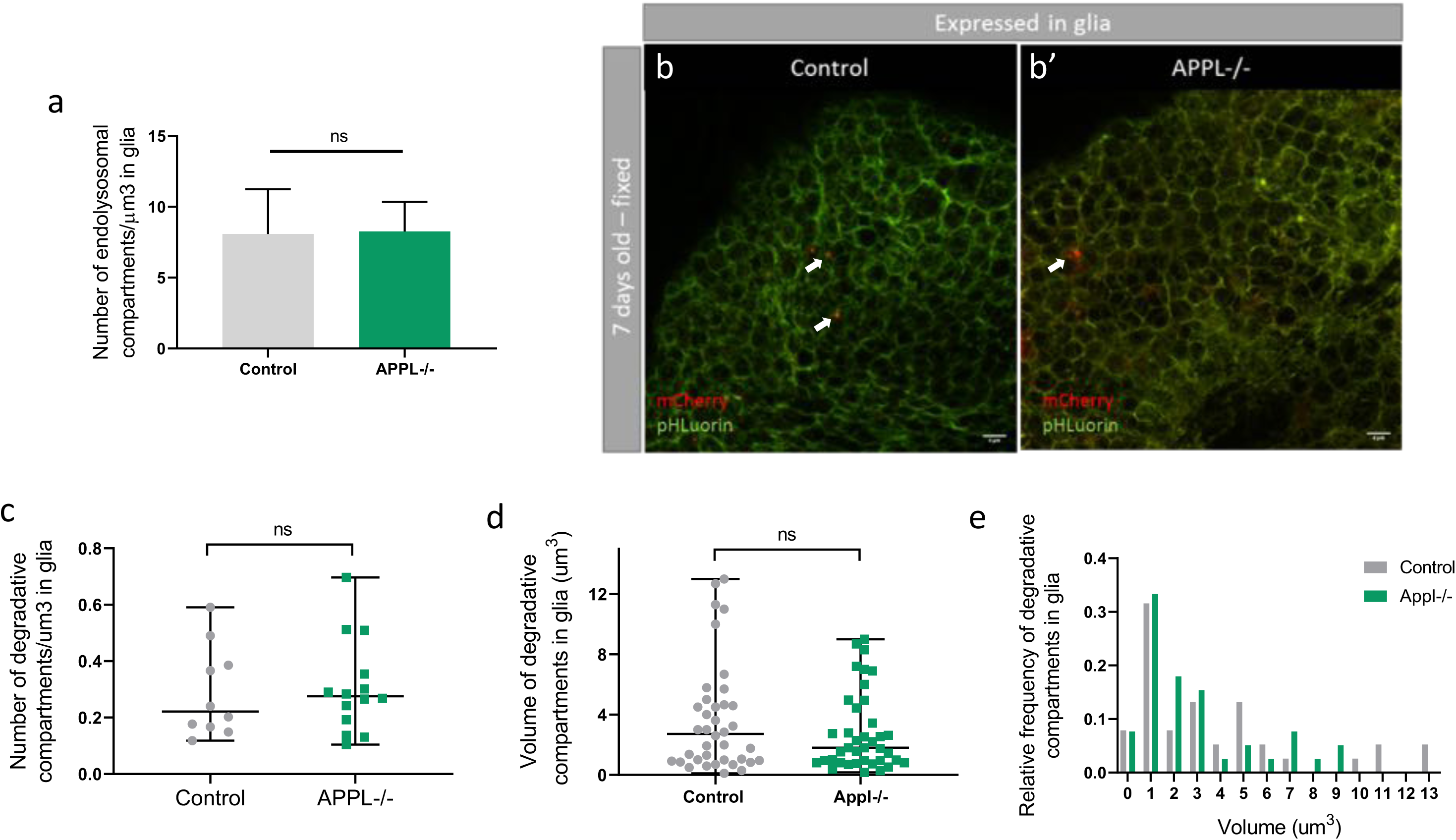
APPL effects on glial endolysosomal network. a, c) The number of endolysosomal and degradative/acidic compartments were not affected by the absence of APPL. b,b’) These confocal slices represent the same area of glial cells but this time from a fixed tissue of control and APPL-/- 7 days old fly brains. d,e) The volume of degradative/acidic compartments was also similar between both conditions.

**Figure S6.**
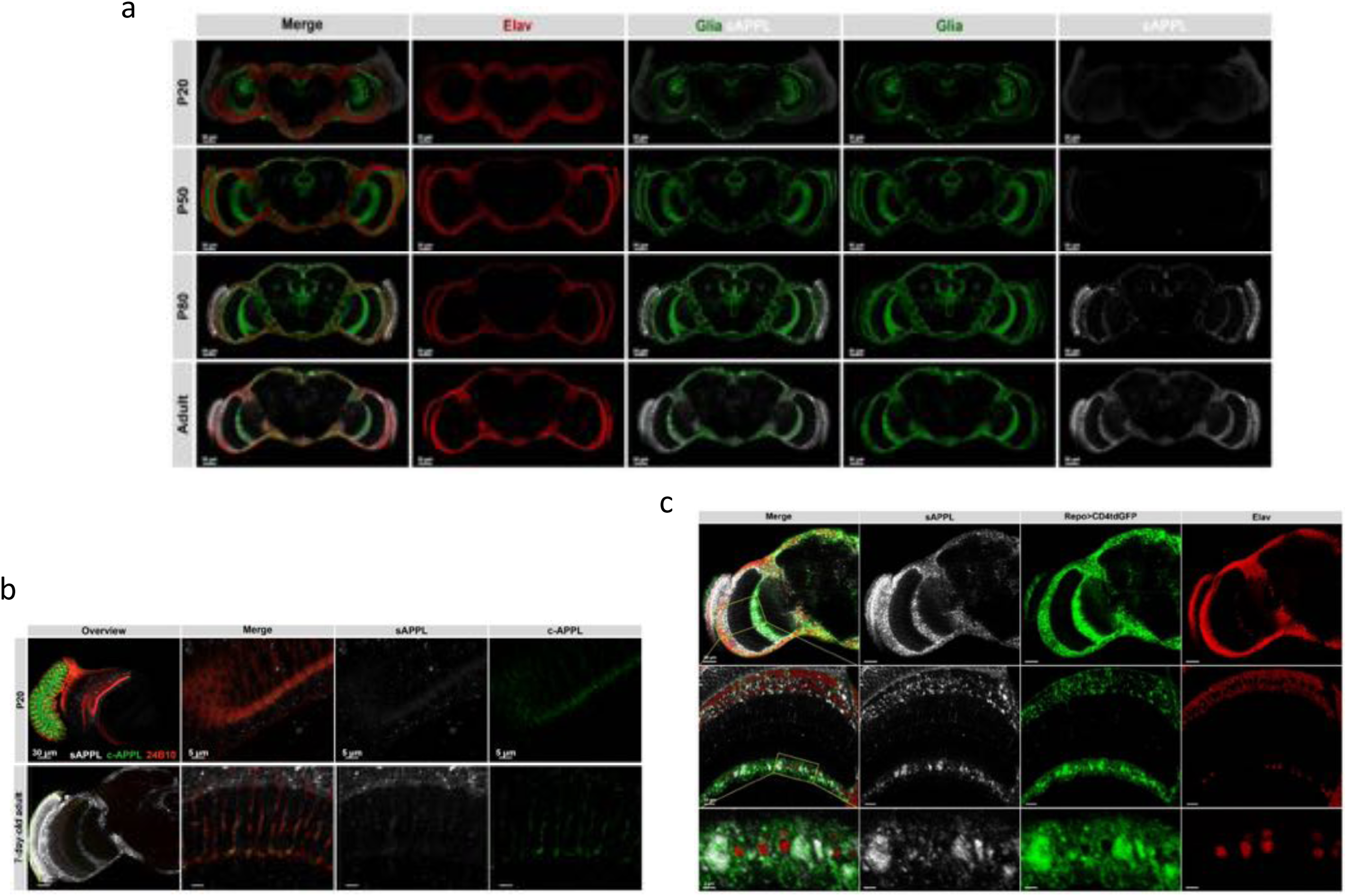
sAPPL travels ubiquitously regardless the site of expression of APPL. a) Confocal sections of a control fly brain throughout development until adulthood that expresses the double fluorescent APPL construct specifically in the optic lobes using the GMR Gal4 driver, ; *UAS-mCherry-APPL-GFP/lexAop-CD4tdGFP; GMR-Gal4/Repo-lexA*. As we can see between P50 and P80 there is a significant release of SAPPL (white) beyond the site of expression reaching all areas of the brain. b) These close-ups on the photoreceptors of the same flies confirm that it is only the SAPPL that travels ubiquitously in the brain, although the C-terminus of APPL, the intracellular part, remains in the cell bodies where it is being expressed. c) This graph highlights that the SAPPL, not only travels throughout the brain, but it also co-localises specifically with the glial marker, Repo (green).

**Figure S7.**
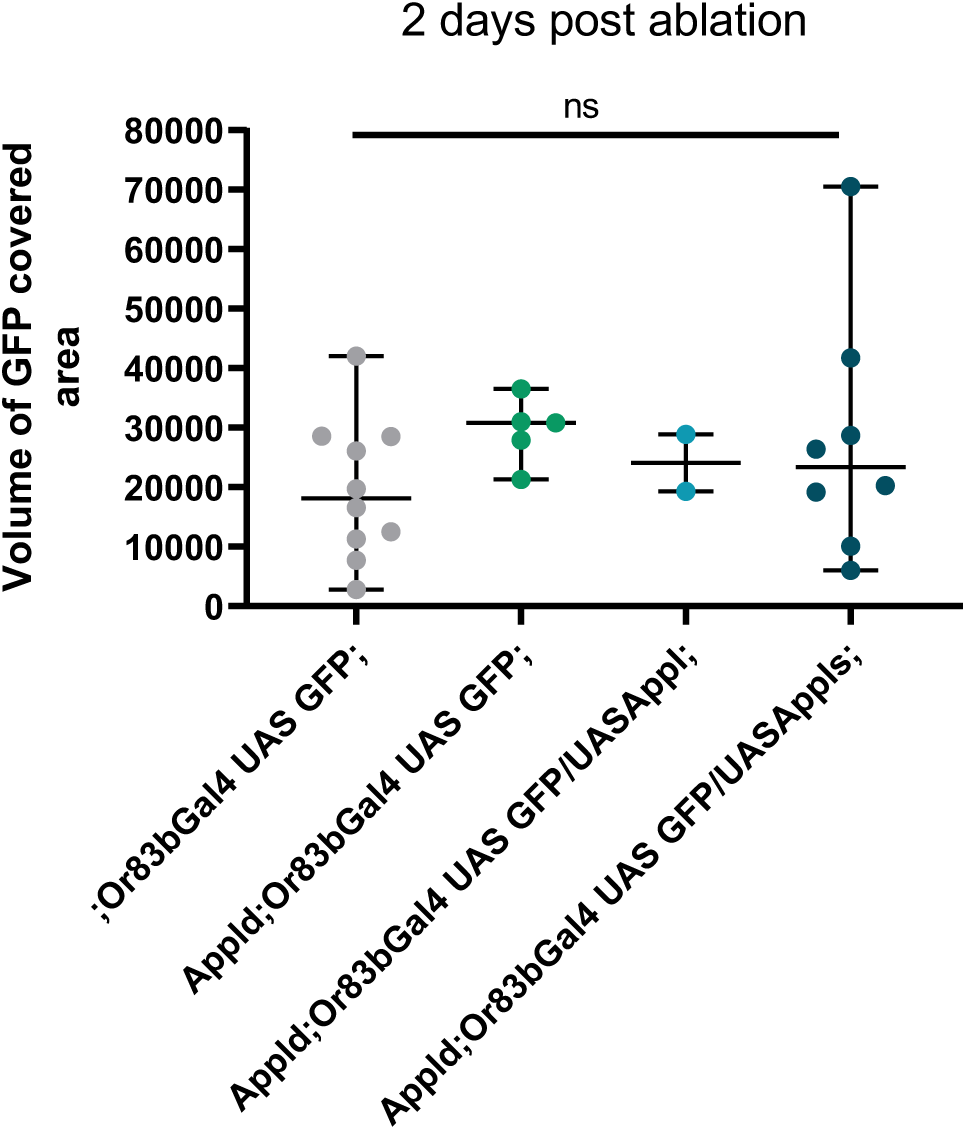
Glial clearance of axonal debris at 2 days post ablation. Quantification of volume of GFP covered region (um3) in the OR83b innervating glomeruli at 2 days post ablation, in control, APPL-/- and the rescue flies: *Appldw*; UAS APPL/OR83bGal4GFP;* and *Appldw*; UAS APPLS/OR83bGal4GFP;.*

